# Automated Retinal Dysplasia Segmentation in Mouse Optical Coherence Tomography Scans Using a UNet-Based model

**DOI:** 10.64898/2026.06.04.730047

**Authors:** Apostolos V. Mikroulis, Miles J. Raishbrook, Marcela Palkova, Jiri Lindovsky, Jan Prochazka, Radislav Sedlacek, Vendula Novosadova, Daniel Novak

## Abstract

**Purpose:** Optical coherence tomography (OCT) is the state-of-the-art non-invasive imaging technique for preclinical retinopathy studies. However, manual pathology annotations in mouse OCT scans are labour-intensive and susceptible to inter-rater variability. To alleviate these issues, we developed a neural network-based model for automated annotation of retinal dysplasia in mouse OCT scans.

**Methods:** Our model was trained on 205 expert-annotated OCT stacks and validated on 40 unseen stacks with additional expert annotation comparisons in a subset of them.

**Results:** The model detects pathologies with high accuracy (F1-score > 0.95) and consistency with experts (median Dice score > 0.8). We integrated the model into a cross-platform app (“OCTOPUS”) that includes batch processing, automated annotation, manual annotation editing, and quantitative longitudinal tracking with pathological area estimation on the fundus image, and exportable results in CSV and SVG formats.

**Conclusions:** Our open-source tool aims to facilitate and standardise OCT assessments between laboratories for efficient preclinical screening and high-throughput phenotyping in mouse models of retinal disease.

## Introduction

Optical coherence tomography (OCT) is the gold standard for non-invasive longitudinal retinal imaging of human retinopathy. OCT provides high-resolution visualisation of retinal structure, neuronal layering, and pathological alterations in murine disease models, supporting phenotypical evaluation and preclinical studies.^1–3^ However, due to the lack of reliable and freely available automated methods for retinal layer segmentation and pathology annotation, manual assessment remains the norm – a process that is labour-intensive, introduces inter-observer variability ^4^ and requires significant expertise.

Most published automated methods focus on retinal layer segmentation to compare mice possessing or lacking a disease state.^5–9^ Although retinal layer thickness is an important metric for evaluating retinopathy, these tools accelerate analysis in longitudinal studies ^10^ but may not capture all pathology-specific aspects of a phenotype. Machine learning (ML) approaches for automated annotation of human OCT datasets are advancing,^11–13^ however there is still a scarcity of tools tailored towards mouse OCT datasets or rapid automated pathology-centric annotation.

## Materials and methods

### Animals

Animals used in this study were wild-type and mutant C57Bl/6N mice of both sexes, bred at the Czech Centre for Phenogenomics (Institute of Molecular Genetics of CAS, Prague, Czech Republic) under standard housing conditions (12/12hr light/dark cycle, food and water *ad libitum*. Optical coherence tomography (OCT) measurements were carried out under general anaesthesia (i.m. Tiletamine + Zolazepam, 30 + 30 mg/kg and 3.2 mg/kg Xylazine) and were kept on a heating pad throughout anaesthesia. Pupils were dilated with 0.5% Atropine (Ursapharm, Prague, Czech Republic) and a transparent gel (Vidisic, Bausch&Lomb, Prague, Czech Republic) was applied to eyes to prevent dehydration. Experiments complied with the ARRIVE 2.0 guidelines, and were approved by the Animal Care and Use Committee of the Czech Academy of Sciences under project license AVCR 6144/2022 SOV II.

### OCT acquisition

Images of posterior eye/retina were acquired by spectral domain–OCT with a standard 30° lens (SD-OCT—Heidelberg-Engineering, Heidelberg, Germany) and an additional +25dpt lens. Anesthetised mice were fitted with +100 dpt contact lenses (Roland Consult, Brandenburg, Germany) and placed on heated platform with the eye to be measured perpendicular to the camera. The retina and fundus were brought into focus, the optic nerve head was centred and volumetric scans consisting of 61 evenly spaced B-scans (cross-sections) per eye were acquired with 77µm spacing between adjacent B-scans. Animals were measured at 20, 28, 36, 44, 52 and 60 weeks of age. The built-in active eye-tracking system and confocal scanning laser ophthalmoscope were used to perform consecutive scans at the identical retinal locations during each measurement.

### Pathology annotation

AVI files exported by Spectralis (Figure 1A) were converted to PNG frames and loaded into the VGG Image Annotator (VIA),^14^ where experienced OCT examiners from the Czech Centre for Phenogenomics manually delineated pathological regions using rectangle bounding boxes. Coordinates were exported to CSV and JSON formats.

**Figure 1.**
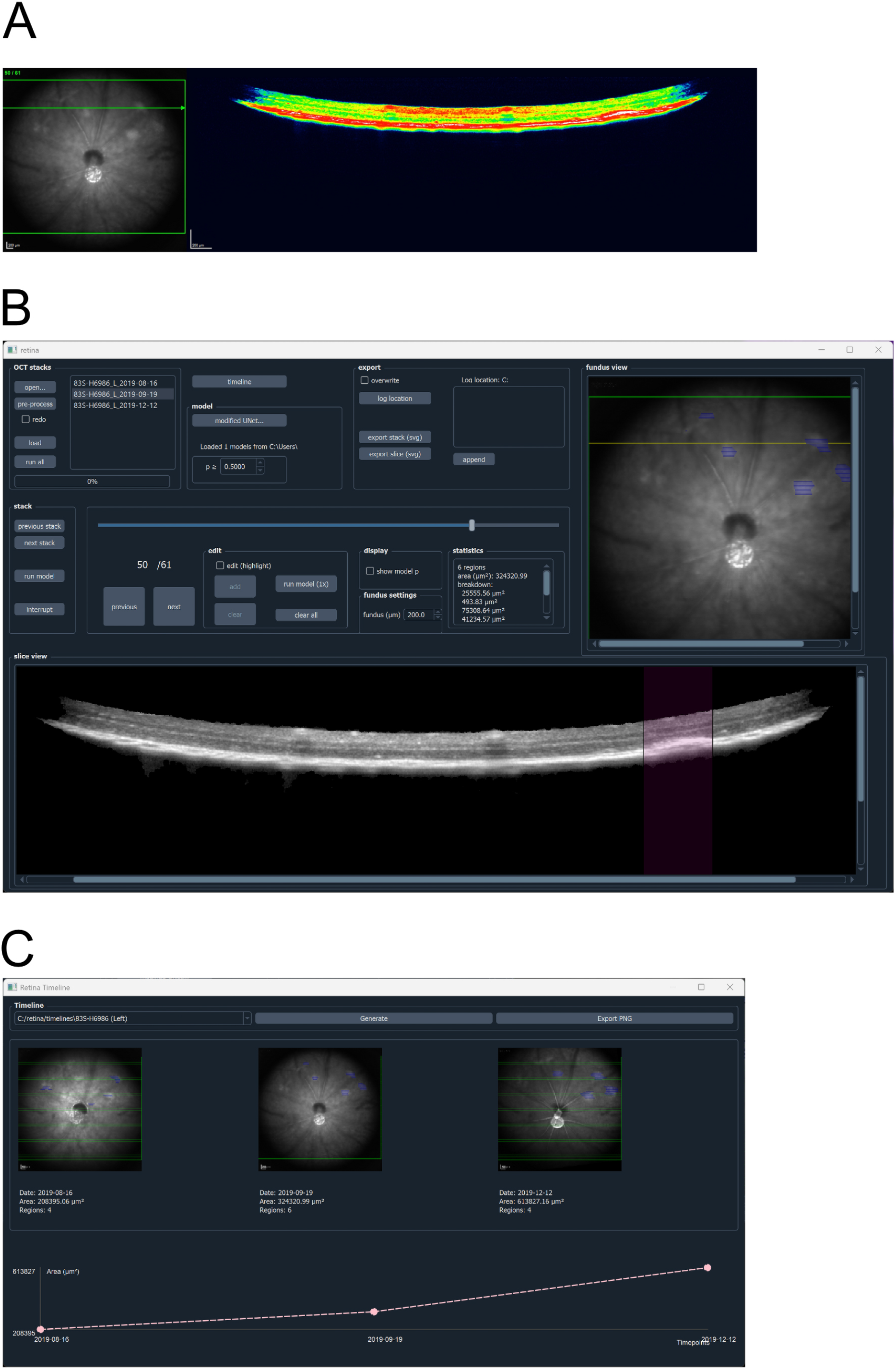
Example video frame and user interface. **(A)** Example of a video frame exported by the Spectralis software. The green overlays on the fundus image (left) indicate the scanning window (green rectangular frame) and the current B-scan (green arrow) depicted on the right. The OCT scan (right) is using a custom jet-like colourmap for greyscale (intensity) values. **(B)** Graphical user interface of the tool. Main window, allowing navigation between and within OCT stacks, model prediction, manual editing of annotations and basic result export functionality. A model-predicted pathological region is displayed in the single slice view panel (light pink shading), and reflected in the fundus view panel (light blue annotation) for the indicated B-scan (yellow line). **(C)** A timeline feature to compare OCT stacks for the same subject at different time points (pathology progression).

### Dataset

The dataset consisted of optical coherence tomography (OCT) images of mouse retinas with localised dysplasia. Pathology annotations performed by experienced OCT examiners from the Czech Centre for Phenogenomics were used as labels for training the machine learning models. A total of n=245 OCT stacks from n = 38 mice were used for the model training and testing. A subset of n=40 OCT stacks was randomly selected at the beginning and set aside for a secondary evaluation of the model. Thus, the model had access to only 205 out of 245 OCT stacks for training and cross-validation.

### Video frame extraction

Frame components (fundus image, scan frame overlay, OCT) were extracted as NumPy arrays. Green overlay lines on the fundus image were restored by mean interpolation from adjacent pixels, and the OCT scan was top-cropped to 400 pixels.

### OCT image preprocessing

OCT images were converted from a custom colourmap to greyscale using a nearest-neighbour mapping derived from paired greyscale and false-colour exports, cropped to 250×1536 pixels, and zero-padded where narrower than 1536 pixels.

The retina region was isolated with the Chan-Vese method with default Scikit-Image parameters,^15,16^ followed by binary morphological post-processing (opening, padding, SDF hole-filling, and erosion) to produce a clean retina mask.

### Modified UNet

The modified UNet ^17^ utilises the preprocessed images. It consists of five down-sampling layers (consecutively doubling sizes from 32 to 512), a middle layer (with size 512), and five up-sampling layers, split with skip connections. The model was constructed using the PyTorch package and exported in ONNX format.

The AdamW optimiser ^18^ was used for the training epochs, with weight decay of 0.05. A learning rate scheduler was used (OneCycleLR ^19^), with learning rate between 2·10^-5^ and 0.002, warm-up for 30% of the cycle and cosine annealing strategy (Figure 2C). The model was trained with a 5-fold cross-validation loop, for 800 epochs per fold.

**Figure 2.**
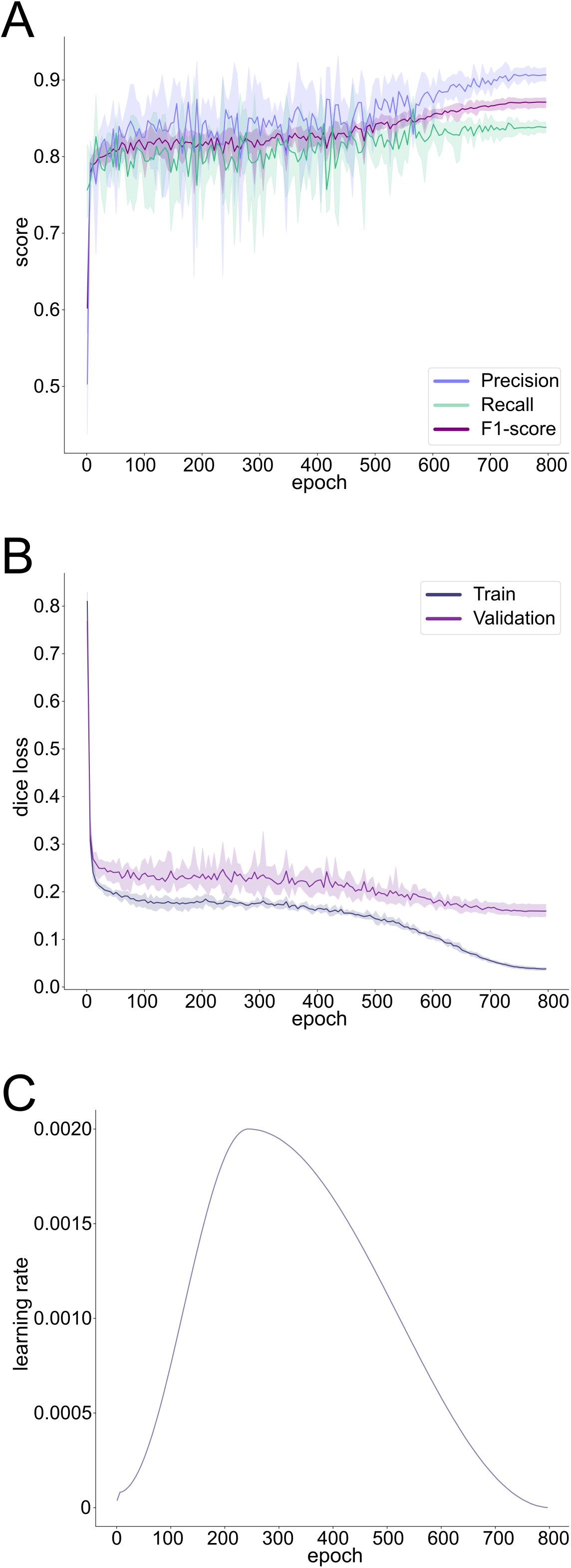
Model training progress. **(A)** Precision, recall and F1-score progress (average and range for 5 cross-validation folds). **(B)** Dice loss for the training sample and validation sample. (average and range for 5 cross-validation folds). **(C)** Learning rate schedule (OneCycleLR).

The dice loss was used (Figure 2B), calculated on pixel columns, and precision, recall and f1-score metrics were tracked for the validation samples, by epoch (Figure 2A). The model was saved for each fold, at the epoch with the highest metrics (highest f1-score) on the validation set.

### Inference

Predicting the pathology segmentation for a new (unseen) image consists of two steps:

(i) extracting and preprocessing the images from the video files, (ii) predicting the segmentation (pixel-column-wise) from the processed image, using the UNet.

### User Interface

The app (“OCTOPUS” – Optical Coherence Tomography Ocular Pathology by U-net Segmentation) uses a graphical user interface (based on the Qt framework) to facilitate the preprocessing and inference process (Figure 1B). The UI supports batch preprocessing and inference across multiple files, and allows navigation and manual editing of annotations within individual OCT stacks. Annotations are reflected in the fundus image panel with semi-automated scale conversion (scale bar size in μm). The GUI tool also estimates the total affected area by interpolating annotated regions between successive OCT slices in the stack and using the fundus image for scaling reference. There is also an option to aggregate annotations (number of annotated regions and their area in μm^2^ on the fundus image) by OCT stack in csv format, and export editable snapshots (layered SVG files) of individual OCT slices and fundus images with their current annotations and overlays. A simple timeline function (Figure 1C) allows quick inspection of pathology progression by tracking the annotated area for longitudinal studies.

## Results

### Training and cross-validation

Training was consistent across all five cross-validation folds, with dice loss and F1-score improving progressively and stabilising in the final 200 epochs, coinciding with the low learning-rate tail of the OneCycleLR schedule (Figure 2).

### Validation on unseen data

The model was also evaluated on new (unseen) data, in comparison with two expert evaluations. The first evaluation was aggregated from the (n=40) folders that were excluded from the model training and their corresponding annotations. The selected models from all five cross-validation folds performed similarly on this dataset (Table 1, mean ± SD across five folds: f1-score 0.954 ± 0.002, ROC AUC 0.973 ± 0.002).

**Table 1.** Summary of validation results for the best model from each cross-validation fold, on unseen OCT stacks (n = 40), compared to expert annotations.

|  | **precision** | **recall** | **f1-score** | **ROC**<br>**AUC** | **Average**<br>**precision** |
| --- | --- | --- | --- | --- | --- |
| *fold 1* | 0.9694 | 0.9419 | 0.9555 | 0.9733 | 0.9377 |
| *fold 2* | 0.9684 | 0.9447 | 0.9564 | 0.9746 | 0.9418 |
| *fold 3* | 0.9650 | 0.9397 | 0.9522 | 0.9715 | 0.9307 |
| *fold 4* | 0.9689 | 0.9379 | 0.9531 | 0.9702 | 0.9331 |
| *fold 5* | 0.9649 | 0.9421 | 0.9534 | 0.9731 | 0.9312 |
| *mean* | 0.9673 | 0.9413 | 0.9541 | 0.9725 | 0.9349 |
| *SD* | 0.0022 | 0.0026 | 0.0018 | 0.0017 | 0.0047 |

The second evaluation, aimed at the consistency of model-expert agreement compared to inter-expert agreement, contained n=32 of the same (unseen) folders, following two additional expert annotations. The model-expert and inter-expert agreement was quantified using the overlap (Dice scores) between annotations (Figure 3): each one of the expert annotations was compared to the other two experts, separately (Figure 3B), and the model predictions were compared to each expert’s annotations (Figure 3C).

**Figure 3.**
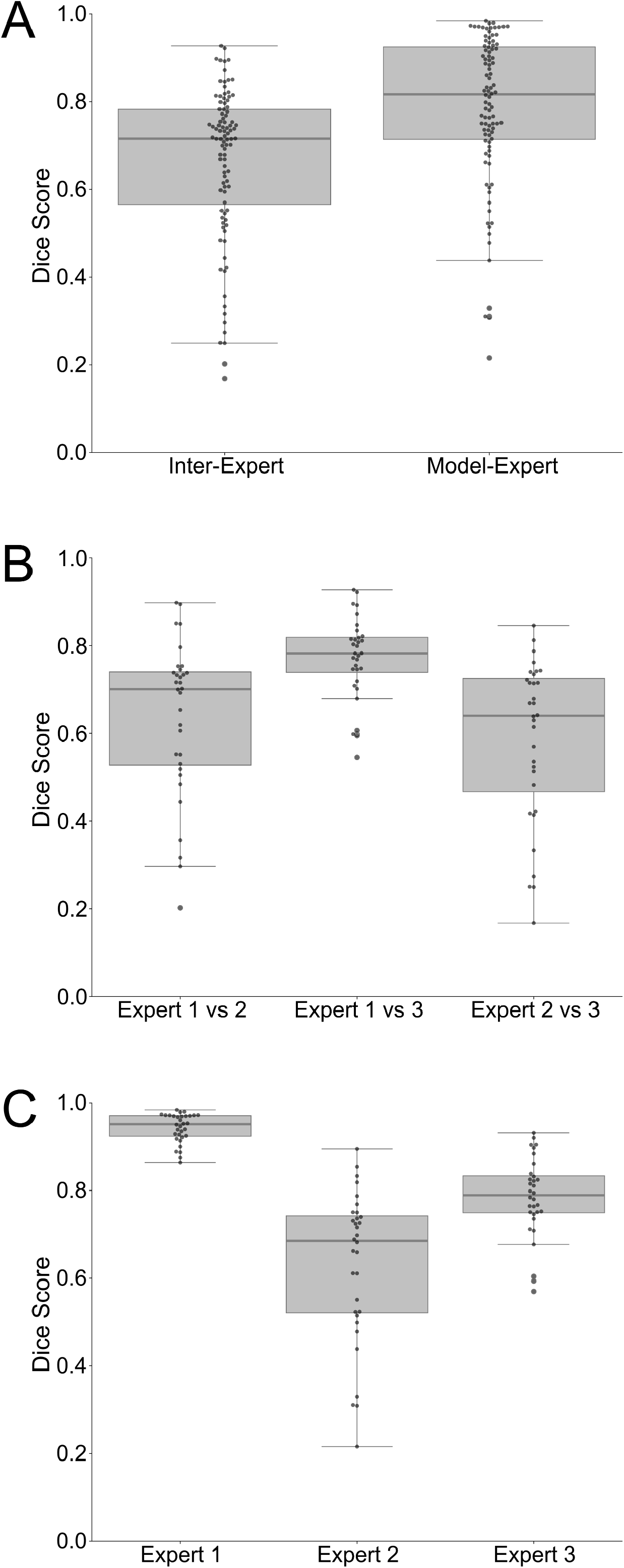
Model and expert annotation consistency between three experts on n=32 OCT stacks novel to the model. **(A)** The model-expert annotation overlap is not inferior to inter-expert annotation overlap. **(B)** Inter-expert annotation overlaps. **(C)** Model annotation prediction overlap with individual experts.

Both inter-expert and model-expert agreement was generally high (median Dice score > 0.7, Figure 3A). Additionally, the model-expert overlap was not inferior to the inter-expert overlap (Figure 3A, one-sided Wilcoxon rank sum test N_1_=N_2_=96, W = 5.130, p > 0.999).

## Discussion

### Comparison with other solutions

Compared to existing tools, like the VGG Image Annotator,^14^ the software tool we developed is more time-efficient (over 10-fold speed-up on large datasets like the one used in the validation we present here), while achieving non-inferior annotation accuracy compared to the manual annotations. In comparison to proposed clinical solutions ^11–13^ and state-of-the-art machine-learning models,^20–24^ we prioritise pathology segmentation instead of pathology classification, since our use case is well-defined pre-clinical retinal dysplasia models – therefore our model focuses on high-accuracy segmentation rather than classification of different pathology phenotypes.

### Limitations

Although our model localises pathological regions with high accuracy, it is also highly specialised in mouse models of retinal dysplasia. As a consequence, different types of pathology (e.g. sparse pathological regions in wild type mice) are not detected sufficiently (f1-score ≈ 0.37 on a small test sample), and we expect similarly low performance in other pathology types without model updates. Additionally, the detection model is tailored to the Heidelberg-Engineering/Spectralis output, with a maximum image width of 1536 pixels. Different manufacturer formats or larger image dimensions will need an additional adapter function or a recompiled model.

### Conclusions

We developed a UNet-based model for automated retinal dysplasia annotations in OCT scans, addressing a gap in preclinical retinal image analysis. Our model performs at high accuracy on unseen data (f1-score > 0.95), and annotates with high consistency with expert annotations (median Dice score > 0.8). We integrated the model into an open-source cross-platform GUI app with batch processing capabilities and options for manual fine-tuning of pathology annotations, aiming to improve accessibility for researchers and facilitate phenotypical assessments in preclinical research.

## Data availability statement

The data that support the findings of this study are available from the corresponding authors upon reasonable request.

Sample raw data (AVI format) are available at: https://octopus.img.cas.cz/ The software developed for this study is available at https://doi.org/10.5281/zenodo.17582639 under the terms of the GNU GPL-3 license.

The model weights are available at https://doi.org/10.5281/zenodo.17866408 under the terms of the CC-BY 4.0 license.

## Acknowledgments

AVM and DN would like to thank the Research Center for Informatics of the Czech Technical University in Prague for the computational infrastructure (RCI Cluster) used in the deep learning model optimisation, training and validation.

AVM would like to thank Denis Baručić (Dept. of Cybernetics, Faculty of Electrical Engineering, CTU Prague) for his help with the deep learning model optimisation and Jiří Anýž (Dept. of Cybernetics, Faculty of Electrical Engineering, CTU Prague) for his input on the method design.

We would like to thank Květoslava Klajblova, who has helped with data measurement and their evaluation.

## Author contributions

Conceptualisation: JL, VN, DN; Data curation: MJR, MP; Formal analysis: AVM; Funding acquisition: RS, DN; Investigation: MJR, MP; Methodology: AVM, MJR, DN; Project administration: VN, DN; Resources: DN; Software: AVM; Supervision: VN, DN; Validation: MJR, MP; Visualisation: AVM; Writing – original draft: AVM, MJR; Writing – review & editing: AVM, MJR, MP, JL, JP, RS, VN, DN

