## Supplementary Figure 1 for "Automated Retinal Dysplasia Segmentation in Mouse Optical Coherence Tomography Scans Using a UNet-Based model"

### Supplementary material

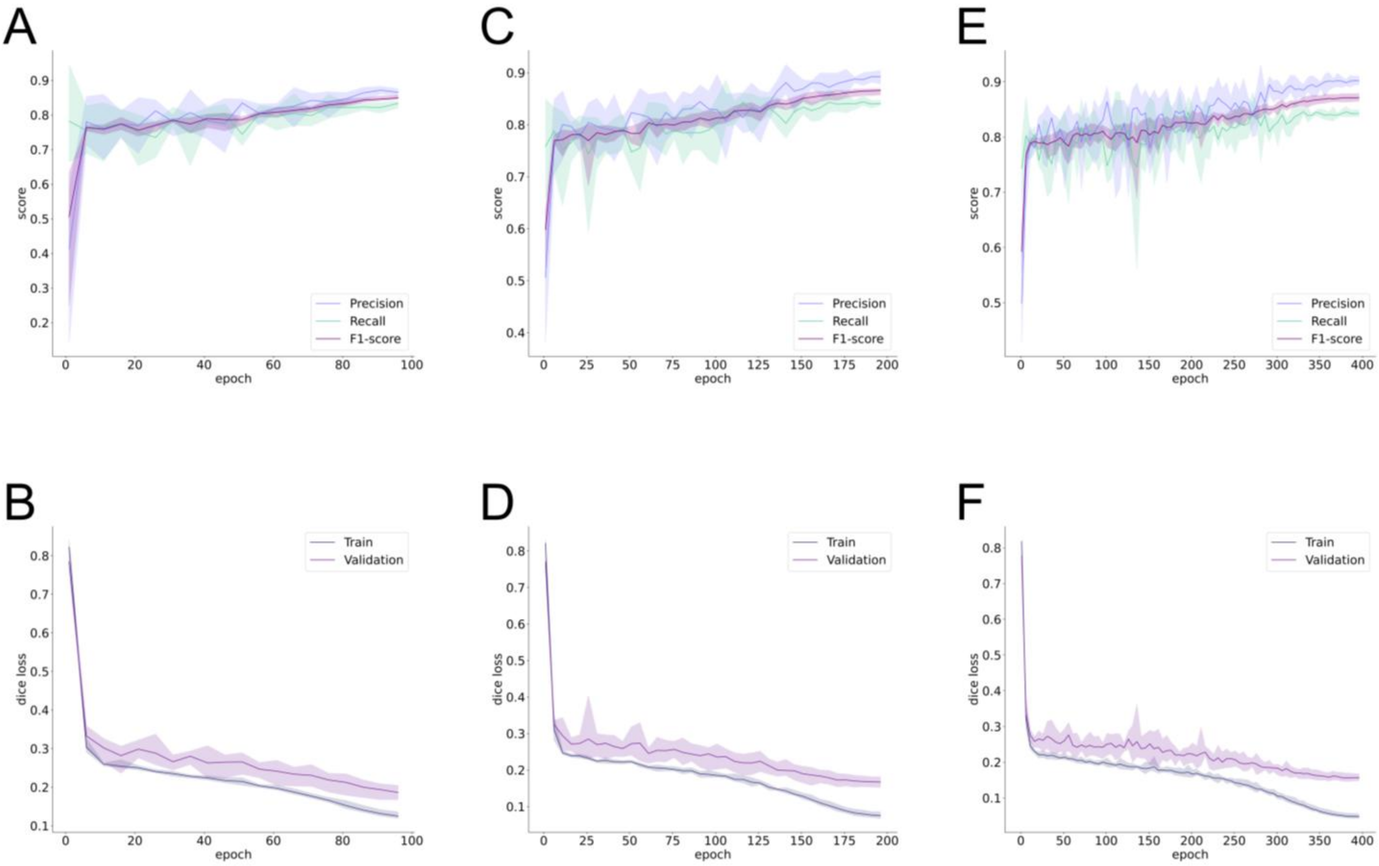


*Figure S 1*. Early model training iterations. **(A-B)** 100 epochs, **(C-D)** 200 epochs, **(E-F)** 400 epochs. **(A, C, E)** Precision, recall and f1-scores calculated on the validation set. **(B, D, F)** Dice loss for the training and validation samples. Averages indicated as solid lines, min—max ranges indicated as shaded areas.
